# Parkinson’s genes orchestrate pyroptosis through selective trafficking of mtDNA to leaky lysosomes

**DOI:** 10.1101/2023.09.11.557213

**Authors:** Mai Nguyen, Jack J Collier, Olesia Ignatenko, Genevieve Morin, Sidong Huang, Michel Desjardins, Heidi M McBride

## Abstract

Inflammation is an age-related factor that underlies numerous human disorders. A key driver of inflammation is the release of mitochondrial DNA (mtDNA), which binds and activates cytosolic sensors. This induces transcriptional responses and, ultimately, pyroptotic cell death. The main challenge has been to understand how mtDNA can cross the two mitochondrial membranes to access the cytosol. Through a genome-wide CRISPR knockout screen we identified a new pyroptotic pathway defined by mtDNA exit within mitochondrial-derived vesicles that are delivered to lysosomes. Critically, breach of lysosomes allows mtDNA to access cytosol, requiring multiple Parkinson’s Disease-related proteins and Gasdermin pores, identified in the screen. These data place mitochondria-to-lysosome transport as a driver of pyroptosis and link multiple PD proteins along a common pathway.

**One sentence summary:** Parkinson’s disease-related proteins regulate pyroptosis

## MAIN TEXT

Cell death occurs in response to external cues that range from developmental signals to inflammation to metabolic stress (*1*). It has long been established that mitochondria act as executioners of multiple cell death pathways, yet the signalling events that couple the external cues to the initiation of cell death from mitochondria remain poorly understood. It is known that these signals induce alterations in mitochondrial architecture, including cristae remodeling, stabilization of ER contact sites, and fragmentation. The **M**itochondrial **A**nchored **P**rotein **L**igase, MAPL, is a SUMO E3 ligase that has a key role in the morphological transformation of mitochondria during apoptotic cell death (*2*). Mice lacking MAPL develop spontaneous hepatocellular carcinoma due, in part, to the strong protection against genotoxic stress-mediated apoptosis in liver, which is a key driver of cancer progression (*3*). In the brain, selective loss of MAPL in parvalbumin interneurons rendered newborn pups fully protected against anesthesia-induced cell death and memory loss (*4*).

While MAPL deletion protects against apoptosis, ectopic expression of MAPL drives cell death (*3, 5*). To characterize the molecular mechanisms by which MAPL promotes cell death pathways, we infected human hepatocellular carcinoma HUH7 cells with adenovirus to express FLAG-tagged MAPL (Ad-MAPL) alongside two controls: reverse tetracycline transactivator (Ad-rtTA), or a MAPL deletion mutant lacking the RING domain (Ad-ΔRING) required for cytosolic SUMOylation activity (*6*). Whereas rtTA did not affect cell viability, exogenous MAPL expression induced cell death and caspase activation in a RING-dependent manner (Fig. 1A, Sup Fig. 1A). Ad-MAPL efficiently targeted mitochondria, as seen previously (*2*) (Sup Fig. 1B). To test whether the ectopic expression of MAPL killed cells in a canonical BAX-dependent apoptotic pathway, we infected baby mouse kidney (BMK) cells, an established cell model where BAX/BAK are deleted, with Ad-MAPL. As a positive control, we used adenovirus to express truncated BID (tBID), which directly activates BAX to drive apoptosis (*7*) (Fig. 1B). Although both tBID and MAPL expression efficiently induced caspase cleavage and cell death in wild type BMK cells, only tBID-induced cell death was fully dependent on BAX/BAK (Fig. 1B, Sup Fig. 1A) (*8*). This indicated that MAPL-induced cell death occurs through a BAX/BAK-independent pathway.

**Figure 1:**
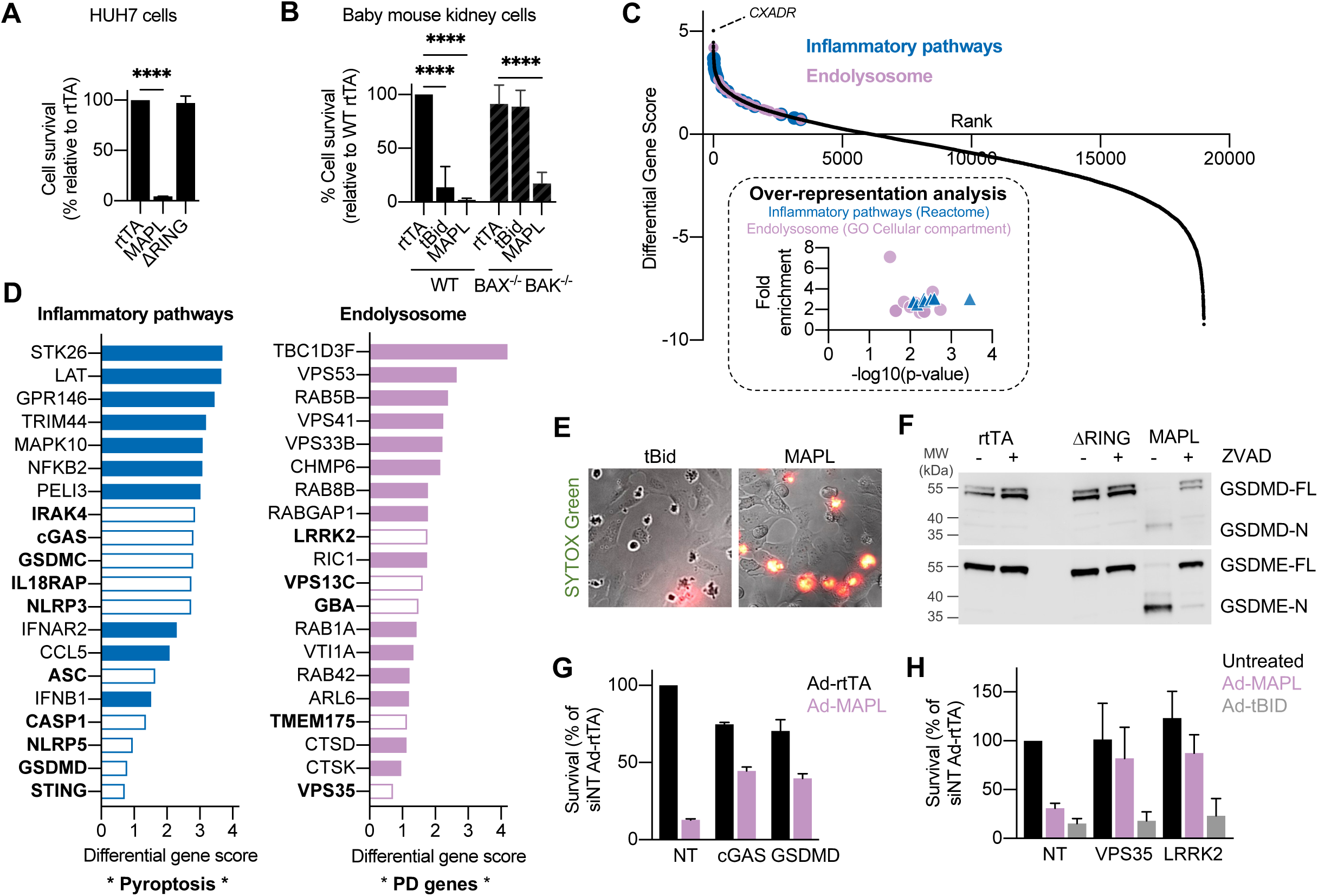
Genome-wide CRISPR knockout screen reveals MAPL induces pyroptosis. (A) Cell viability was measured in HUH-7 cells expressing empty vector (rtTA), MAPL, or MAPL without its RING domain (ι1RING) for 24 hours (N = 6 independent experiments). (B) Cell viability was measured in wild type and BAX/BAK double knockout (DKO) baby mouse kidney cells (BMKs) after expression of rtTA, tBID or MAPL (N =4). (C) Plot of differential gene scores (MAPL - rtTA) mean value from 4 sgRNAs per gene) of the genome wide CRIPSR knockout screen. (D) Plot of differential gene scores. (E) SYTOX Green assay after tBID or MAPL expression. (F) Representative immunoblot full length (-FL) and cleaved (-N) Gasdermins D and E (GSDMD/E) GAPDH was used as a loading control (N = 3). (G) Knockdown of cGAS or GSDMD enhanced cell viability in MAPL-overexpression relative to rtTA. (H) Knockdown of VPS35 or LRRK2 enhanced cell viability in MAPL-but not tBID-overexpression (N = 3). For all graphs, values represent means. Error bars represent SEM. Comparisons were facilitated by One-way ANOVA. ****P<0.0001.

### Genome-wide CRISPR survival screen reveals MAPL induces pyroptosis

To characterize this BAX/BAK-independent death pathway, we performed a genome-wide CRISPR knockout screen to identify genes whose loss confers resistance to MAPL-induced cell death in HUH-7 liver cells. The validity of this approach was internally confirmed given that the top ranked gene in our screen was the coxsackievirus and adenovirus receptor (*CXADR*), essential for infection and MAPL expression (Sup Table 1, Fig. 1C). Gene set enrichment analysis identified genes required for MAPL-induced cell death that were linked to inflammation and indicative of pyroptosis (inset Fig. 1C, and D). Pyroptotic cell death is driven by the cytosolic inflammasome comprised of the sensor (NLRP3), adaptor (ASC) and the effector Caspase-1 proteins (*9*), all of which were identified within the screen. We also identified key substrates of Caspase-1, Gasdermins D and C, whose cleaved N-terminal fragments form pores in the plasma membrane leading to cell rupture, a hallmark feature of pyroptosis (Fig. 1D) (*10*). Additionally, we identified other proteins that can modulate the inflammasome including a cytosolic sensor of double-stranded DNA cGAS (cyclic GMP-AMP synthase encoded by *MB21D1)*, and a downstream of cGAS activator of interferon signalling STING1 (encoded by *TMEM173*) (Fig. 1D) (*11*).

Analysis of enrichment within cellular compartments identified components of endolysosomal membranes as requisite for MAPL-induced cell death (Fig. 1D). This included vesicle tethering subunits of the HOPS (VPS33b) and CORVET (VPS41) complexes (*12*), as well as ESCRT-III machinery responsible for the generation of intralumenal vesicles within the multivesicular body (CHMP6 and CHMP4B) (*13*), suggesting that endolysosomal identity and trafficking are essential for cell death. Notably, the screen also identified multiple Parkinson’s disease (PD)-related genes linked to the endomembrane system, including VPS35, LRRK2, GBA, TMEM175 and VPS13C (Fig. 1D) (*14*), suggesting that at least a subset of PD proteins act along a common pathway in the regulation of pyroptotic cell death.

To validate the major hits in the screen, we first tested whether MAPL overexpression killed cells through pyroptosis. We investigated the hallmark plasma membrane rupture by monitoring cellular uptake of a cell impermeant DNA stain, SYTOX green (*10*). Ectopic expression of MAPL induced SYTOX green uptake, whereas tBID overexpression led to apoptotic blebs with no breach of the plasma membrane, further demonstrating the distinct nature of the apoptotic (tBID) and pyroptotic (MAPL) death pathways (Fig. 1E) (*1*). Western blot analyses confirmed that expression of MAPL, but not ΔRING, led to the cleavage of two caspase substrates Gasdermins D and E, a process required for pyroptosis (Fig. 1F) (*10*). This cleavage was inhibited in the presence of the pan-caspase inhibitor ZVAD, consistent with cleavage by caspase 1 (Fig. 1F). Moreover, silencing Gasdermin D or cGAS protected against MAPL-induced cell death, as did silencing of VPS35 or LRRK2 (Fig. 1G and H, Sup Fig 1). Gasdermin D depletion also blocked MAPL-induced SYTOX green uptake (Sup. 1C). Depletion of these proteins did not affect tBID-induced apoptotic cell death (Fig. 1H), suggesting a unique role for the lysosome and PD-related proteins in pyroptosis. Altogether, these results show that MAPL overexpression induces pyroptotic cell death in a manner dependent on its SUMO ligase activity through a pathway requiring multiple proteins involved in PD and endolysosomal trafficking.

### VPS35 and MIROs drive vesicular removal of mtDNA from mitochondria

The identification and validation of cGAS as an essential component of MAPL-induced pyroptosis suggested that exposure of either nuclear or mitochondrial DNA within the cytosol was essential to drive this cell death pathway (*15*). Immunofluorescent staining of double-stranded DNA revealed a MAPL and RING-dependent increase in the presence of cytosolic DNA foci (Fig. 2A and B). To determine whether these foci originated from mtDNA or nuclear DNA, we passaged U2OS cells with ethidium bromide to generate Rho0 cells depleted of mtDNA, and validated with immunofluorescent imaging (Sup Fig. S2A) (*16*). Cytosolic DNA foci were lost in Rho0 cells expressing MAPL (Sup Fig. 2). Importantly, Rho0 cells were protected against MAPL-induced pyroptosis (Fig. 2C), confirming that mtDNA is required for MAPL-induced pyroptotic death.

**Figure 2:**
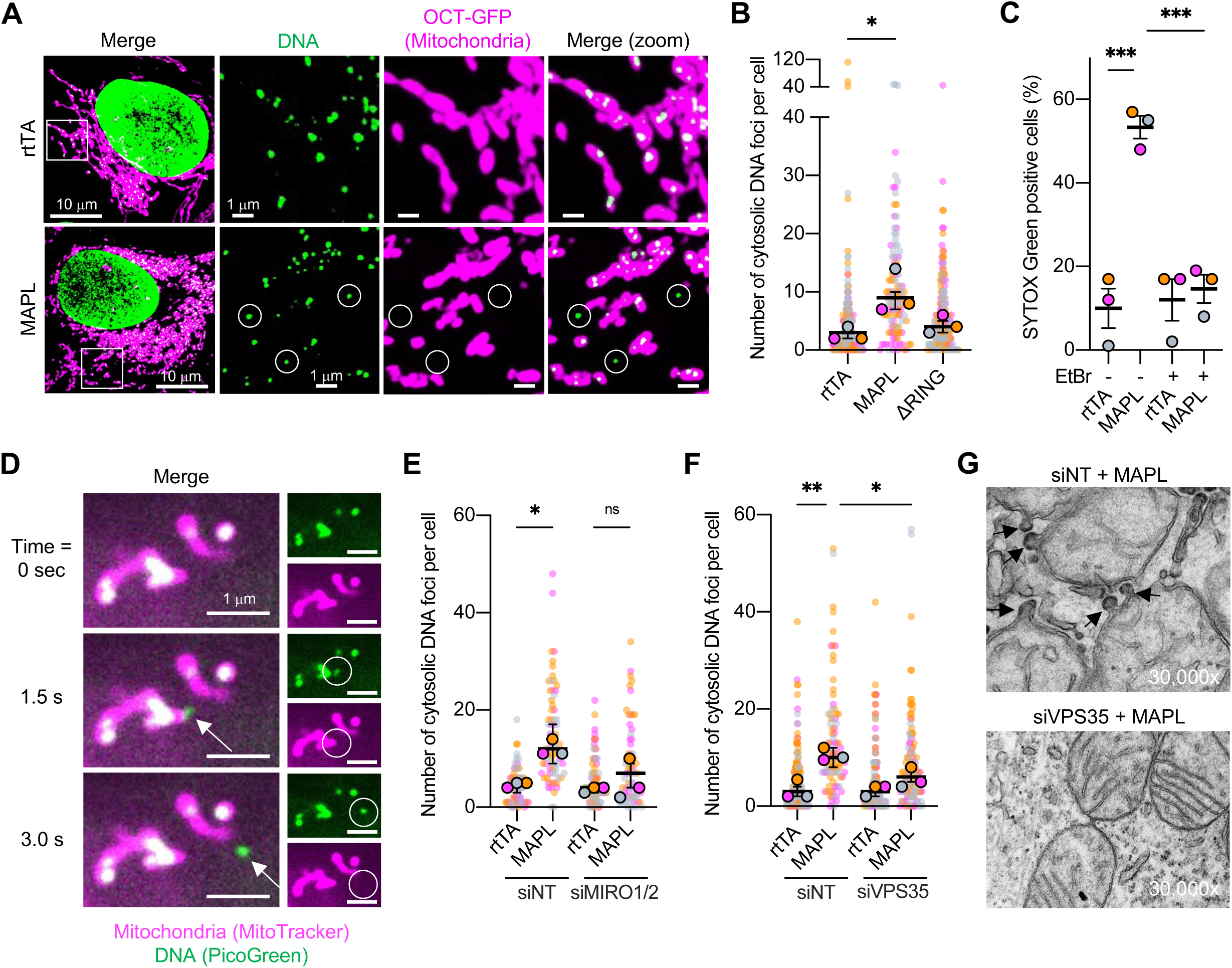
MAPL triggers VPS35- and MIROs-dependent release of mtDNA in mitochondrial-derived vesicles. (A) Representative fluorescent images of U2OS showing mitochondrial network and mitochondrial DNA (mtDNA) following expression of MAPL compared to control rtTA. (B) Quantification of mtDNA release after 48h rtTA, MAPL or DRING expression in U2OS cells stably expressing mitochondrial matrix marker OCT-GFP. (C) Rho0 cells are resistant to MAPL-overexpression SYTOX Green uptake. (D) Time-lapse confocal microscopy showing an mtDNA efflux event from the mitochondrial network. PicoGreen visualises DNA and MitoTracker DeepRed visualises mitochondria. (E) Silencing MIRO1 and MIRO2 reduced the amount of DNA foci in the cytosol following MAPL overexpression. (F) Knockdown of VPS35 significantly reduced cytosolic mtDNA foci. (G) Electron micrographs showing mitochondrial protrusions and MDVs after MAPL expression in U2OS cells treated with non-targetting siRNA (siNT), but not in U2OS cells VPS35-targetting siRNA (siVPS35). Different colours represent independent repeats, with background circles showing corresponding values for individual cells. For graphs in B, E and F, values represent medians and error bars represent 95% CI. Graph C shows mean values +/- SEM. Comparisons were facilitated by One-way ANOVA. *P<0.05, **P<0.01, ***P<0.001.

Next, we sought to understand how mtDNA is released from mitochondria. Two major pathways facilitating the release of mtDNA from mitochondria have been described (*17*). Entire nucleoids can be extracted via inner mitochondrial membrane herniation through large BAX/BAK pores after cytochrome c release (*18*). Alternatively, mtDNA fragments can be released through inner membrane mitochondrial permeability transition pores and outer membrane VDAC pores (*19–21*). Since our cell death pathway is independent of BAX/BAK and the mtDNA foci are too large to pass through VDAC pores (Fig. 2A), we sought alternative mechanisms for mtDNA release in the cytosol. Gasdermin D and E pores have also been shown to facilitate mtDNA release from mitochondria directly (*22–25*), however silencing three gasdermins C, D and E together had no effect on the appearance of cytosolic mtDNA foci (Sup Fig. 2C) while fully protecting against cell death (Fig 1G, Sup Fig 1C). cGAS was also not required for MAPL-induced mtDNA release consistent with both gasdermins and cGAS acting downstream of mitochondria in this pathway (Sup Fig. 2D). Using a DNA intercalating dye PicoGreen and time-lapse imaging, we observed the active and rapid release of nucleoid-sized mtDNA foci from mitochondria with a linear trajectory, reminiscent of mitochondrial derived vesicle (MDV) production (Fig. 2D and Sup. Vid. 1). MDVs are cargo-selective structures formed by the lateral tubulation of mitochondrial membranes along microtubules via the Rho GTPases MIRO1 and MIRO2 before DRP1-mediated scission (*26*). The speed and linear trajectory of the mtDNA movements was consistent with MIRO driven vesicle budding and transport. Indeed, knockdown of MIRO1/MIRO2 significantly reduced the amount mtDNA release into the cytosol after MAPL expression, as seen by confocal imaging (Fig. 2E). These data indicate that mtDNA is released within MDVs upon expression of MAPL.

We previously showed that the PD-related protein VPS35, whose depletion protects against MAPL-induced cell death (Fig. 1H), interacts with MAPL to regulate the trafficking of MDVs to peroxisomes (*27*). Silencing VPS35 robustly decreased the number of cytosolic mtDNA foci (Fig 2F). Ultrastructural analyses revealed that MAPL overexpression led to a striking generation of electron dense double-membrane protrusions from mitochondria (Fig. 2G). Critically, these protrusions were lost upon VPS35 silencing (Fig. 2G). These data identified the PD related protein VPS35 as a regulator of vesicular mtDNA release, essential for pyroptotic cell death.

### mtDNA is trafficked to endolysosomal subcompartments via MDVs

MDVs are known to deliver mitochondrial cargo to endolysosomal compartments (*28–30*). Consistent with this, ectopic expression of MAPL led to a significant increase in mtDNA within LAMP1-positive endolysosomes (Fig. 3A and B). Live imaging also showed PicoGreen DNA-positivity in a subset of dextran-labelled lysosomes in cells expressing MAPL (Fig. 3C and D). Quantification of this subset of lysosomal structures showed mtDNA positivity in ∼15% of lysosomes within the cell (Fig. 3D and E). Time lapse imaging captured long-range transport of MDVs to lysosomes already containing mtDNA (Fig. 3F, Sup. Vid. 2), supporting the notion that this subset of lysosomes is specifically targeted by MDVs. We also observed lysosomal arrival proximal to the mtDNA exit sites, enabling direct and rapid transfer of content from mitochondria to lysosomes. (Fig. 3G, Supp. Vid. 3). These data identify lysosomes as the target organelle for mtDNA-containing MDVs.

**Figure 3:**
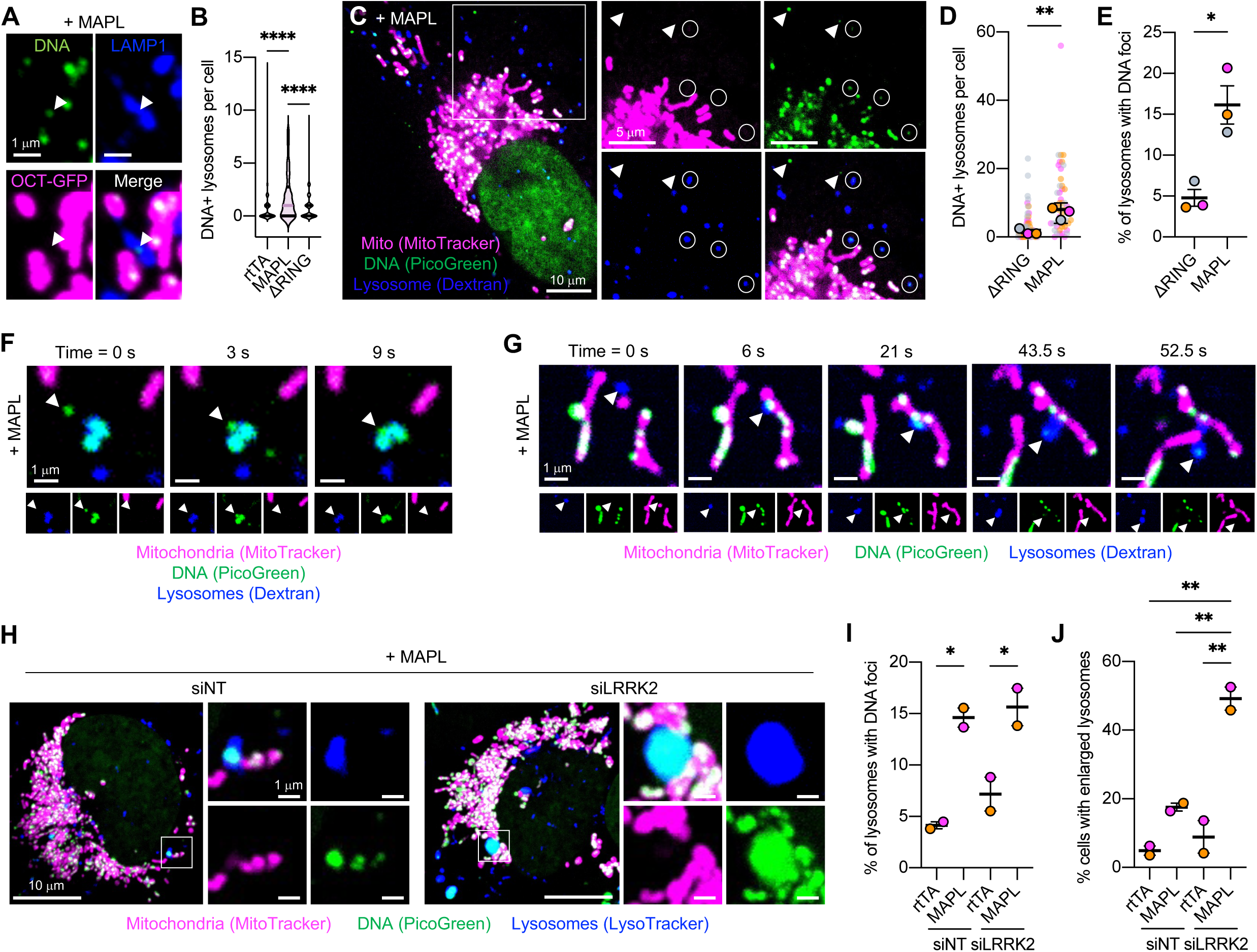
mtDNA is trafficked to endolysosomes. (A) U2OS-OCT-GFP cells overexpressing MAPL for 48 hours were immunostained with DNA and LAMP1. Arrowhead show an example of mtDNA-bound LAMP1-positive endolysosome. (B) Quantification of the number of mtDNA-bound LAMP1+ endolysosomes per cell after expression of rtTA, MAPL or DRING for 48h in U2OS-OCT-GFP. (C) Live cell imaging showing the presence of cytosolic mtDNA (circles) and endolysosome-bound (arrowheads) mtDNA after 24 hours of MAPL expression. (D) Quantification of DNA+ Dextran+ endolysosomes per cell and (E) percentage of Dextran+ endolysosomes that contain DNA. Time-lapse confocal microscopy showing (F) the delivery of cytosolic mtDNA to an endolysosome and (G) the recruitment of an endolysosome to the site of mtDNA efflux and immediate sequestration of the mtDNA-bound MDV. (H) Enlarged lysosomes appear upon MAPL overexpression in si-LRRK2-treated cells but not si-NT-treated cells. (I) Quantification of DNA in LysoTracker+ endolysosomes and (J) endolysosomal enlargement in cells expressing rtTA or MAPL after LRRK2 silencing. Different colours represent independent repeats, with background circles showing corresponding values for individual cells. For graph D, values represent medians and error bars represent 95% CI. Graphs in E, I and J show mean values +/- SEM. Comparisons were facilitated by One-way ANOVA. *P<0.05, **P<0.01, ****P<0.0001.

Given the reported roles for LRRK2 in the regulation of lysosomal function (*31–35*) we next examined the potential requirement for LRRK2, identified in our CRISPR screen, in the delivery of mtDNA+ MDVs to the lysosomes. The data reveal that mtDNA is successfully transported and delivered to lysosomes in MAPL-expressing cells lacking LRRK2 (Fig 3H, I). Notably, we observed the appearance of a subset of lysosomes (∼1-5 per cell) that enlarged into vacuolar like structures in LRRK2 silenced cells upon expression of MAPL (Fig 3H, quantified in Fig 3J). This is indicative of lysosomal stress, a phenotype seen in LRRK^-/-^ tissues that generated similar enlarged, vacuolar lysosomes upon treatment with chloroquine *in vivo* (*33*). We next sought to determine how LRRK2 silencing protect cells from MAPL-induced killing while it does not affect mtDNA delivery to the lysosomes.

### mtDNA release from leaky lysosomes depends upon gasdermins and LRRK2

With mtDNA within the lysosome, the question of how it is released to cytosol for the activation of cGAS remained unsolved. It has been shown that conditions of lysosomal stress can cause a breach in the membrane leading to content release (*36*), which could include mtDNA. Ruptured lysosomes can be monitored through the recruitment of cytosolic β-galactoside-binding lectin proteins (galectins), that bind exposed glycan-modified proteins normally protected within an intact endomembrane system (*36*). Recruitment of Galectin 3 to LAMP1-stained lysosomes was seen in MAPL-expressing cells (Fig. 4A). Similar to cell death and mtDNA release, recruitment of GAL3 to ruptured lysosomes was dependent on MAPL ligase activity (Fig. 4B). Importantly, the GAL3 positive lysosomes were enriched on the population that specifically labelled with PicoGreen in a time dependent manner (Fig. 4C [circles] and D). Ultimately, about 45% of GAL3+ lysosomes were positive for mtDNA (Fig 4E). Given that only 15% of all lysosomes contain mtDNA (Fig 3D), this indicates an enrichment and selectivity of GAL3 recruitment to mtDNA containing structures. We hypothesized that the overexpression of GAL3 would selectively repair lysosomal membrane wounds and promote cell survival (*36*). Indeed, transient overexpression of GAL3-RFP within MAPL-infected cells led to increased survival in a colony formation assay (Fig. 4F). This provides evidence that lysosomal breach is a key step of the death pathway.

**Figure 4:**
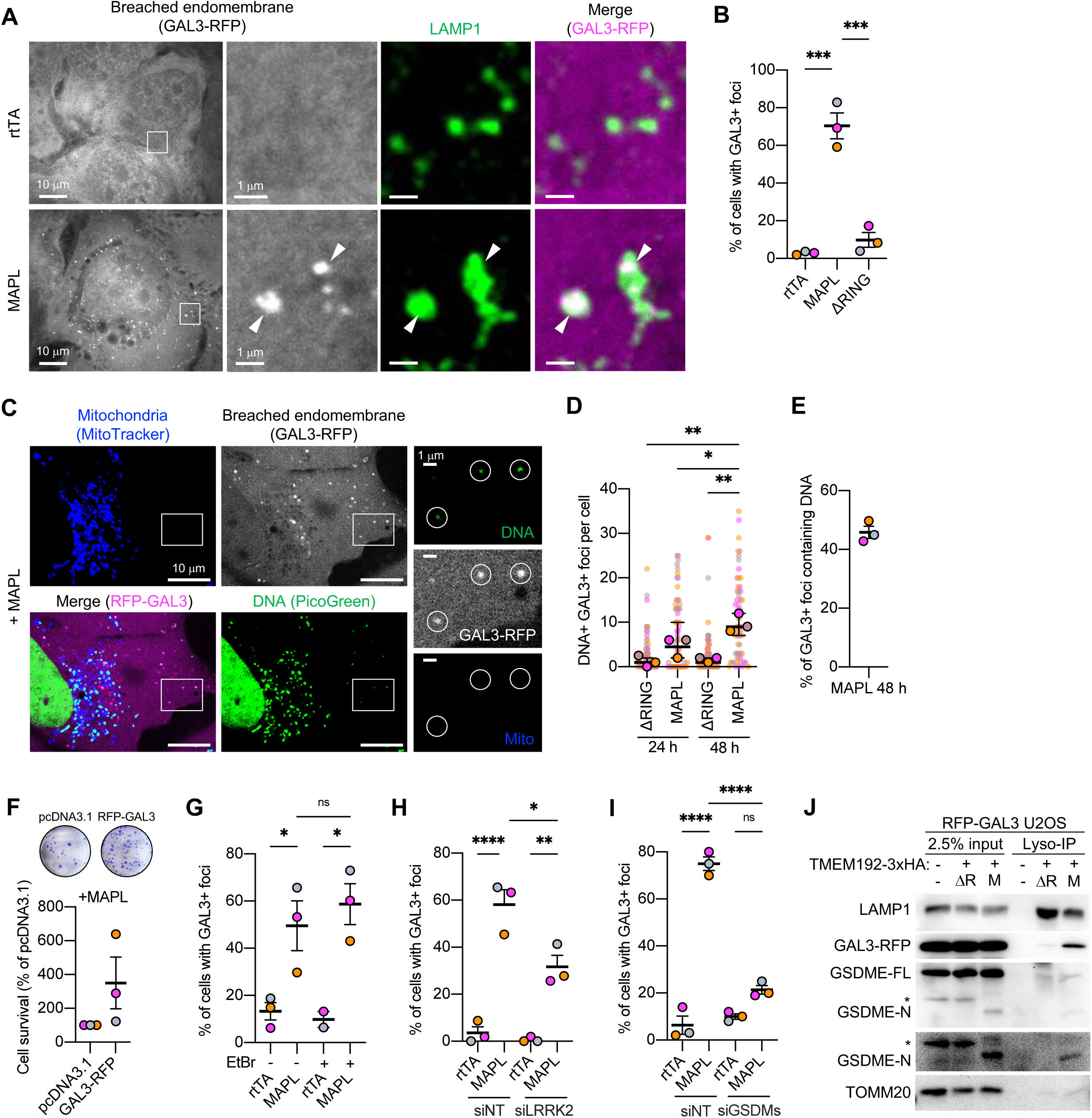
Breach of endolysosomes exposes mtDNA to cytosolic DNA sensors. (A) Representative image of U2OS cells stably expressing RFP-Galectin-3 (U2OS-RFP-GAL3) showing GAL3 recruitment to LAMP1 positive structures (arrows). (B) Quantification of endolysosomal damage following the overexpression of rtTA, MAPL or Δ1RING. Cells with extensive lysosomal damage (>10 GAL3 foci per cell) are considered GAL3 positive cells. (C) Live imaging showing mtDNA in GAL3-positive endolysosomes after MAPL expression. (D) The number of mtDNA- and GAL3-positive foci are induced by MAPL expression in a time dependent manner. (E) Percentage of GAL3-positive foci containing DNA. (F) Transient expression of GAL3 confers resistant MAPL-induced cytotoxicity measured by colony formation. (G) Quantification of endolysosome breach in wild type (parental) and Rho0 143b cells after rtTA or MAPL expression. (H) Quantification of MAPL-induced endolysosome breach after knockdown of GSDMC, GSDMD and GSDME (siGSDM) compared to non-targetting siRNA treatment (siNT). (J) Immuno-isolation of lysosomes from U2OS cells expressing TMEM192-3XHA. Western blotting showed N-terminal fragments of Gasdermin E, and GAL3 in isolated lysosomes from MAPL expressing cells. (*) nonspecific band. Different colours represent independent repeats, with background circles showing corresponding values for individual cells. For graph D, values represent medians and error bars represent 95% CI. Graphs in B, E, F, G, H and I show mean values +/- SEM. Comparisons were facilitated by One-way ANOVA. *P<0.05, **P<0.01, ***P<0.001, ****P<0.0001.

We next wondered what caused lysosomal breach upon expression of MAPL. The recruitment of GAL3 observed preferentially on mtDNA-containing lysosomes suggested that DNA itself may signal lysosome breach. However, MAPL-induced recruitment of GAL3 to lysosomes was unaffected in Rho0 cells lacking mtDNA (Fig. 4G). This indicates that DNA was not required to initiate lysosomal breach, and that lysosomal breach alone is insufficient to drive cell death, given that Rho0 cells are resistant to pyroptosis (Fig 2C). We next examined screen hits linked previously to membrane integrity, including LRRK2 (*22, 32, 34*) and gasdermins (*10, 37*). While mtDNA was delivered to lysosomes in LRRK2 silenced cells (Fig 3H, I), there was a significant reduction in GAL3 recruitment onto lysosomes in MAPL expressing cells (Fig. 4H). This could reflect a direct requirement for LRRK2 in mediating recruitment of GAL3 to broken lysosomes (*34*), in which case there would be increased damage and DNA leakage. Alternatively, the lysosomes may have remained intact, explaining why GAL3 is not recruited. Given that loss of LRRK2, like overexpression of GAL3, is protective against MAPL induced death, it suggests that the lysosomes remain intact.

Like siLRRK2, silencing gasdermins D and E also resulted in a near complete absence of GAL3 recruitment to lysosomes (Fig. 4I). To biochemically test whether the pore-forming N-terminus of gasdermins may have assembled directly within breached lysosomes, we generated stable U2OS cells stably expressing a lysosomal membrane marker TMEM192-3xHA in order to immunoisolate lysosomes (*38*), and GAL3-RFP. Western blotting revealed that the cleaved N-terminal fragment of endogenous gasdermin E, along with GAL3-RFP, were present in lysosome fractions isolated from MAPL-expressing cells in a RING-dependent manner (Fig. 4J). This demonstrates that gasdermin pores form selectively on mtDNA-containing lysosomes.

### MAPL regulates inflammatory responses in primary macrophages

Of note, the screen identified a series of kinases and regulators of the TLR4 and MyD88 inflammatory pathways (Fig 1D). It has previously been shown that TLR4 activation via lipopolysaccharide (LPS), which mimics gram negative bacterial infection, induces the release of mtDNA into the cytosol, where it can activate the NLRP3 inflammasome (*21, 39–41*). We first tested whether LPS promoted mtDNA release in foci in U2OS cells. Indeed, we observed mtDNA release, similar to those observed following MAPL expression, within 3-6 hours of LPS exposure in U2OS cells (Fig. 5A and B). Time-lapse confocal microscopy captured further evidence that lysosomes are recruited to sites of mtDNA efflux in MDVs under inflammatory conditions (Fig. 5C). In the example shown, a mtDNA-containing MDV remained in contact with a motile lysosome over a 33 second time frame, and appeared to be within the lysosome at 46 seconds, suggesting a tethering event prior to fusion or microautophagy. We then tested the role of MAPL in a professional immune cell and treated primary bone marrow derived macrophages (BMDM) with LPS and interferon-ψ (IFN-ψ). We observed mtDNA release (Fig. 5D) and delivery to lysosomes (Sup Fig. 3). Importantly, mtDNA release was not induced in BMDMs derived from MAPL KO mice, confirming the importance of MAPL in driving mtDNA release (Fig 5E). In addition, LPS/IFNg-induced pyroptotic cell death was robustly inhibited in the absence of MAPL (Fig. 5F). Together, these data demonstrate the commonality of the MAPL-induced pyroptotic pathway identified in our CRISPR screen with the canonical TLR4 signalling induced with LPS treatment. This places MAPL and Parkinson’s-related proteins as a key regulator of inflammatory signals.

**Figure 5:**
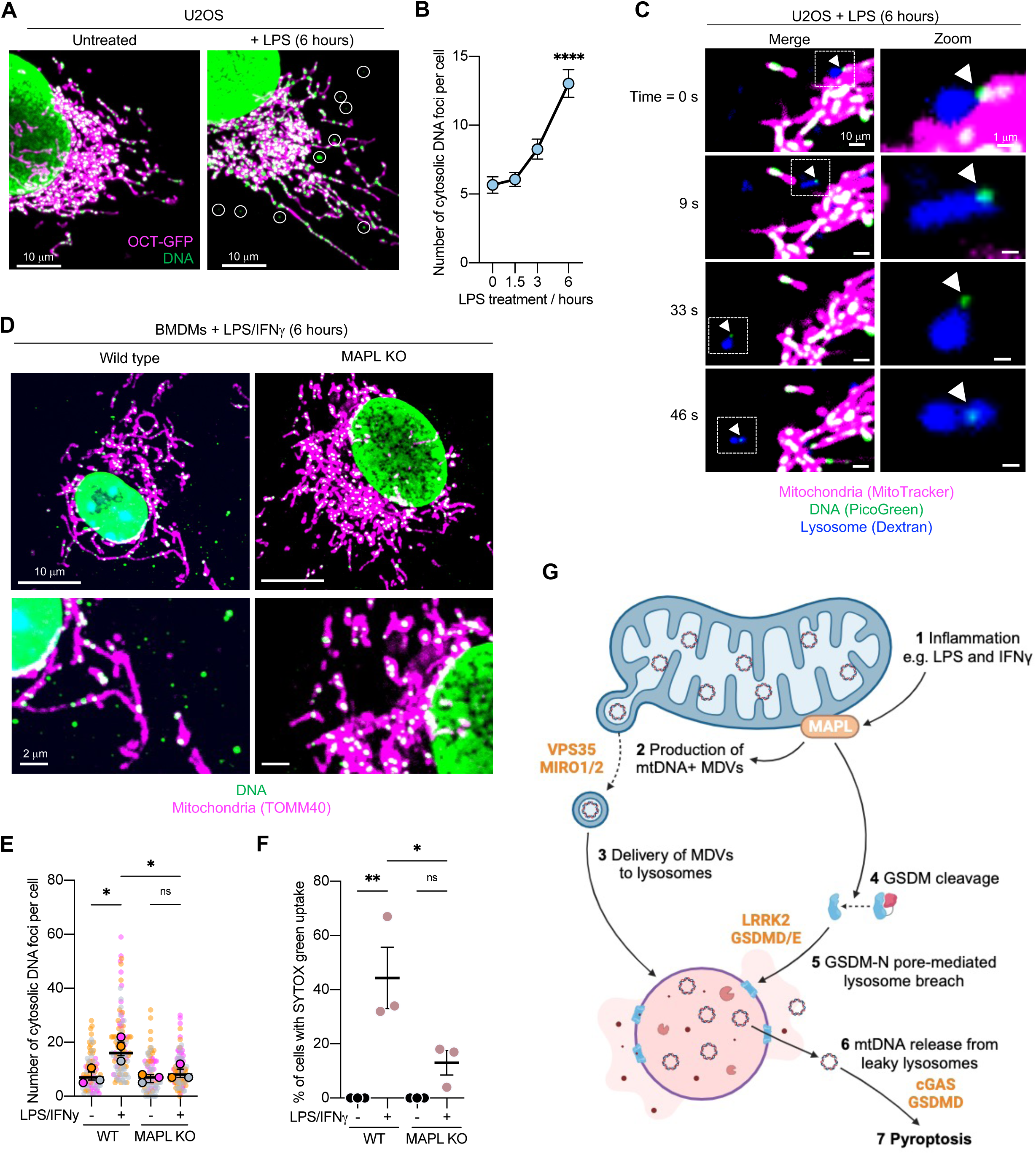
MAPL regulates responses to physiological inflammation. (A-B) Lipopolysaccharide (LPS) induced mtDNA release in U2OS-OCT-GFP in a time dependent manner. Representative images from 6 hours are shown in A, with circles highlighting cytosolic DNA+ foci. (C) Time-lapse confocal imaging showing recruitment of an endolysosome to the site of MDV-mediated mtDNA efflux prior to its trafficking and delivery. (D-E) Lipopolysaccharide (LPS) and interferon-gamma (IFNψ) treatment of wild type (WT) bone marrow-derived macrophages (BMDMs) for 6 hours resulted in the release of mtDNA in a MAPL-dependent manner. (F) Measurement of pyroptotic cell death by SYTOX Green uptake in WT and MAPL KO BMDMs following LPS-IFNψ treatment for 36 hours. (F) An overview of a model for MAPL-induced inflammatory cell death.

## DISCUSSION

The mechanisms by which mtDNA is released into cytosol is of profound importance given its association with many human disorders from cancer to neurodegeneration and infectious diseases(*11*). The discovery that “leaky lysosomes” act as a means of mtDNA release into cytosol provides new insight into the engagement of the cGAS/STING pathway downstream of MDV mediated transport to lysosomes (Fig 5G). mtDNA was seen recently to exit mitochondria within MDVs in a cancer model where cGAS/STING were activated (*42*). However, the mechanisms allowing DNA release into cytosol from the MDV remained unresolved and we suspect may involve transit through the lysosomes. Similarly, our mechanisms may be important in the age-associated mtDNA release and STING activation observed recently in mouse models of neurodegeneration, senescence, inflammation, and aging (*43*).

In seeking the mechanisms of MAPL-induced cell death we unite multiple PD-related proteins, including VPS35 and LRRK2, along a common pyroptotic pathway. VPS35, a known interactor of MAPL that regulates MDV transport (*27*), was critical for the formation of mtDNA containing MDVs upon MAPL overexpression. This finding may provide a mechanistic basis for recent observations of VPS35-dependent mtDNA delivery to endosomes (*44*). While VPS35 acts at mitochondria, loss of LRRK2 led to alterations within the lysosome. The delivery of mtDNA to a subset of lysosomes that recruit Gal3, suggested that this unique lysosome population was susceptible to Gasdermin pore insertion. *In vitro* studies have demonstrated that the insertion and assembly of N-terminal gasdermin fragments shows a preference for specific lipid compositions, including cardiolipin and phosphatidylserine (*45, 46*). We consider that depletion of LRRK2, GBA, VPS13C, and TMEM175, which play key roles in lysosomal lipid flux and metabolism (*47–50*), would result sorting deficits that would result in an inability to generate this specific compartment. The loss of other screen hits within the endomembrane family further supports this idea, including sorting complexes (HOPS, CORVERT and ESCRT machinery) (*12, 13*).

Altogether, using the power of an unbiased genome-wide CRISPR screen we place the mitochondria-lysosome axis at the heart of pyroptotic cell death involving mtDNA trafficking and release. Our work provides a new framework to study the underlying contribution of mtDNA release and activation of cGAS/STING in a host of pathological settings from aging to Parkinson’s disease, cancer and inflammation.

## Supporting information

Sup. Vid. 1

Sup. Vid. 2

Supp. Vid. 1

## ACKNOWLEDGEMENTS

We thank all members of the Desjardins ASAP team and members of the lab for critical comments throughout the course of this study.

## AUTHOR CONTRIBUTIONS

Conceptualization: MN, MD, HMM

Methodology: MN, JJC, OI, GM, SH, HMM

Investigation: MN, JJC, OI, GM

Visualization: MN, JJC, OI, GM, HMM

Funding acquisition: HMM

Project administration: HMM Supervision: HMM

Writing – original draft: JJC, HMM

Writing – review & editing: MN, JJC, OI, GM, SH, MD, HMM

## Data and materials availability

Materials are available upon request. All data are available in the main text or the supplementary materials.

## METHODS

### Ethics Approval

Animal experimentation was conducted in accordance with the guidelines of the Canadian Council for Animal Care. Protocols were approved by the Animal Care Committees of McGill University.

### Culturing of cell lines

HUH-7 and U2OS cells were cultured in DMEM containing 4.5 g/L glucose, L-glutamine, and sodium pyruvate (Wisent, 319-027-CL), supplemented with non-essential amino acids (NEAA; Gibco, 11140050) and 10% fetal bovine serum (FBS; Wisent, 085-150). Cell lines were cultured in a humidified incubator at 37°C and 5% CO_2_ and regularly tested for mycoplasma using the MycoAlert mycoplasma detection kit (Lonza, LT07-418).

### Isolation of bone-marrow derived macrophages (BMDMs)

MAPL KO mice were generated and maintained as previously reported (*3*). Mice were from a C57BL/6J background, registered as RRID:MGI:7524577. To prepare BMDMs, marrow was flushed from mouse tibias and femurs using 25-gauge needle with DMEM. The resulting homogenate was filtered through 40 μm mesh filters. Cells were collected by centrifugation and plated on uncoated culture dishes in DMEM, 10% heat inactivated FBS and 25% L929 cell conditioned supernatant. The resulting macrophages were collected after 8 days in culture for experiments. BMDM were routinely treated with 1.5 μg/ml of LPS (Invitrogen, tlrl-3pelps) and 0.2 μg/ml IFNγ (Peprotech, 315-05) for the indicated time.

### Transfection and generation of stable cell lines

U2OS cells were transfected with 2.5 μg of plasmid per well (6 well plate) at approximately 80% confluence using Lipofectamine 2000 (ThermoFisher, 11668019) according to manufacturer’s instructions. The following plasmids were used: pmRFP-C1-Galectin-3 (a kind gift from Maximiliano Gutierrez (*34*)) pOCT-GFP (*51*), and TMEM192-3xHA (RRID:Addgene_102930). To generate stable cell lines, two days after transfection cells were sparsely plated in 150 mm dishes in media containing 750 μg/ml G418 for two weeks. Single clones were subsequently isolated. Confirmation of ectopic gene expression was determined by confocal microscopy and/or immunoblot.

### RNA interference

siRNA-mediated knockdown of genes was facilitated by Lipofectamine RNAiMAX (Invitrogen, 13778150) according to manufacturer’s instructions and confirmed by immunoblotting. siRNA was used at a final concentration of 20 nM. The following human On-TARGET plus individual or Smart Pool siRNAs (Dharmacon) were used: cGAS/MB21D1 (L-015607-02), GSDMC (L-014716-02), GSDMD (L-016207-00), GSDME/DFNA5 (L-011844-00), LRRK2 (L-006323-00), MIRO1 (L-010365-01), MIRO2 (L-008340-01) and VPS35 (J-010894-05).

### Cells lysates, PAGE and immunoblotting

Cells were solubilised in lysis buffer comprising 50 mM HEPES pH7.4, 150 mM NaCl, 1% Triton X-100, 1 mM EDTA followed by centrifugation for 10 minutes at 10,000*g* and 4°C. Total protein concentration was determined using Bradford assay (Bio-Rad) before samples were mixed with sample buffer (50 mM Tris pH6.8, 0.1% glycerol, 2% SDS, 100 mM β-mercaptoethanol) and heated to 95°C for 5 minutes. Protein extract was separated with SDS-PAGE using a 4%-20% gradient gel, before transfer to 0.22 μm pore supported nitrocellulose membrane (Bio-Rad). Western-Lightning PLUS-ECL (Perkin-Elmer) with an INTAS ChemoCam (INTAS Science Imaging GmbH) was used to visualise bands, before processing with ImageJ software.

The following primary antibodies and dilutions were used: AIF (Santa Cruz, sc-13116, 1:1000), cGAS (Cell signaling, 15102, 1:1000), GBA (R&D Systems, MAB410-SP, 1:1000), GSDMD (Abcam, ab210070, 1:1000), GSDME (Abcam, ab215191, 1:1000), LRRK2 (Abcam, ab133474, 1:1000), MAPL (Sigma, HPA017681, 1:1000), VPS35 (Abnova, H00055737, 1:1000), RHOT1 (Sigma, HPA010687, 1:1000), Vinculin (Sigma, V4505, 1:1000).

### Cell death and pyroptosis assays

Cell survival was determined using Promega CellTiter-Glow Luminescence Cell Viability Assay kit as described by the manufacture. Briefly, cells were plated in triplicates in 96-well culture dish and cultured overnight (1 x 10^5^ cells per well). Cells were infected with adenovirus vectors (20 pfu/cell) for 2 days after which time CellTiter-Glow reagent was added and luminescence was determined using Promega GlowMax luminometer.

Pyroptotic cell death was determined via SYTOX Green uptake. Monolayer cells plated on glass bottom dishes were treated for indicated time followed by addition of SYTOX green at final concentration of 1 μM directly in the medium and immediately imaged. Images were acquired using Zeiss Axio Observer.Z.

### Preparation of adenovirus and infection of cells

Adenovirus expressing MAPL and MAPL-delta RING were custom generated for us by Vector Biosystems (*3*). Ad-tBID was a gift from Dr. Gordon Charles Shore, McGill University. The virus was expanded by one round of infection of Ad-293 cells (Agilent 240085) at 5 plaque forming units (PFU) per cell in complete growing media for 3-5 days. Cells were collected and lysed by 3 rounds of freeze/thaw. Viral titre (plague forming unit, PFU) was determined by plaque forming assay. Briefly, serial dilution of the viral stock was used to infect Ad-293 for 2 hours after which time 1% low-melting agarose was overlaid. After 2-3 weeks, visible plaques were counted to determine PFU.ml^-1^. For transient expression, cells were routinely incubated with adenovirus encoding rtTA, MAPL or ΔRING at 20-30 PFU per cell in complete media. Expression was confirmed by immunoblotting and/or immunofluorescence.

### Immunofluorescence and image acquisition

For immunofluorescent microscopy, cells were grown on uncoated glass coverslips (ThermoFisher, 12-545-81). After treatment, media was removed from coverslips and cells were directly fixed with 5% (wt.vol^-1^) paraformaldehyde (PFA; Sigma, P6148) in PBS (prewarmed to 37°C) for 15 minutes at 37°C. Coverslips were washed three times in PBS, before 10-minute quenching with 50 mM NH_4_Cl. Cells were washed again three times in PBS followed by permeabilization using 0.1% Triton-X100 in PBS for 10 minutes. After three washes in PBS, cells were blocked with 5% FBS in PBS for 10 minutes, then sequentially incubated in primary antibody (prepared in 5% FBS in PBS) for 1 hour. The following primary antibodies and dilutions were used: DNA/mtDNA (Abcam, ab27156, 1:1000), LAMP1 (Cell signaling, 9091, 1:500), TOMM40 (Proteintech, 18409-1-AP, 1:1000). Between primary antibodies, there were three washes in PBS for five minutes each. After the final primary antibody and washes, cells were incubated in the appropriate secondary antibodies (1:2000) for 1 hour: cross-adsorbed goat anti-mouse IgG Alexa Fluor 488 (A28175) or 568 (Invitrogen, A11004), and cross-adsorbed goat anti-rabbit IgG Alexa Fluor 647 (Invitrogen, A21245). Cells were then washed twice in dH_2_O, before addition of DAPI (Invitrogen, D1306) in dH_2_O for 10 minutes to stain nuclei. After one further wash in dH_2_O, coverslips were mounted onto microscope slides (VWR, 82003-412) with approximately 10 μL of mounting media (Dako, S302380-2).

Images were acquired using an Olympus IX83 confocal microscope containing a spinning disk system (Andor/Yokogawa CSU-X) using an Olympus Olympus UPLANSAPO Å∼100/1.40 numerical aperture (NA) oil objective and Andor Neo sCMOS camera and MetaMorph software. Laser lines corresponding to 405 nm, 488 nm, 555 nm, and 637 nm were used, with excitation and exposure times maintained across samples for each independent experiment. Z-stacks of 0.2 μm per stack were used.

### Live cell microscopy

Cells plated on uncoated glass-bottom dishes (MatTek, P35G-1.5-20C, size no. 1.5) then treated as described. For co-staining mtDNA and the mitochondrial network, cells were first washed three times using 1X PBS, before addition of complete media and incubated with 2.5 μl.ml^-1^ Quant-iT PicoGreen (ThermoFisher, P7589) for 30 minutes. Cells were then washed three times in complete media, then incubated in 10 nM MitoTracker Deep Red FM (Invitrogen M22526) for 30 minutes before imaging. For labelling late endosomes and lysosomes, cells were incubated in 100 nM LysoTracker Red FM for 30 minutes at the same time as MitoTracker staining. For labelling lysosomes, cells were incubated for 6 hours with 0.1 mg.ml^-1^ Dextran Alex Fluor 555, 10,000 MW (Anionic, fixable) then chased for 18 hours in fresh media. Immediately before imaging, HEPES buffer was added to a final concentration of 25 mM. Images were acquired using an Olympus IX83 confocal microscope containing a spinning disk system (Andor/Yokogawa CSU-X) using an Olympus Olympus UPLANSAPO Å∼100/1.40 numerical aperture (NA) oil objective and Andor Neo sCMOS camera and MetaMorph software. Cells were imaging at 37°C in a humidified chamber controlled by an INU TOKAI HIT system. For videos, a rate of 1 frame per 1.5 seconds was used. Acquisition parameters were controlled to limit bleedthrough.

### Image processing and quantification

Images were processed using FIJI Image J software. For counting the number of cytosolic mtDNA foci, a maximum intensity projection was generated, before automated macros enabled unbiased quantification. First, a mask of the nucleus (DAPI signal), mitochondria (TOMM40 or OCT-GFP signal), and mtDNA (DNA signal) were generated. mtDNA foci outside of mitochondria were calculated by subtracting the nuclear and mitochondrial masks from the mtDNA mask, leaving only non-nuclear and non-mitochondrial foci. Cell perimeter was drawn by hand for each cell, and foci within the perimeter were counted. The number of mDNA+ lysosomes was counted by eye, after manually thresholding the lysosome and DNA channels. For these experiments, the mitochondrial network was also taken into consideration to prevent false positive results. A single z-plane was taken into consideration for these experiments to ensure colocalisation. Quantification of lysosomal damage was also done by eye. Cells with extensive lysosomal damage (over 10 galectin 3 foci per cell) were considered GAL3 positive cells.

### Transmission electron microscopy

Cells were plated on glass coverslips and treated with adenovirus (20 PFU/cell) for 24 hours. Cells were fixed with 2.5% glutaraldehyde, 0.1 M sodium cacodylate, 0.1% CaCl_2_ for 1-2 days at 4°C. Coverslips were treated with 1% OSO_4_, 1.5% ferrocyanide for 45 minutes at 40°C. Following 3 washes with H_2_0, coverslips were incubated with 5% uranyl acetate for 45 min. Coverslips were then washed extensively with ethanol and acetone before embedding in Spurr epoxy resin. Samples were sliced and mounted on grids by McGill university Facility for Electron Microscopy Research (FEMR) and images were acquired with FEI Tecnai G2 Spirit Twin 120 kV Cryo-TEM.

### Pooled genome-wide CRISPR knockout screen

Functional genetic screens were performed in HUH-7 cells using the human CRISPR Brunello lentiviral pooled library (4 unique sgRNAs (single guide RNAs) per gene, comprising a total of ∼76,000 sgRNAs) (*52*). HUH-7 cells (8 x 10^6^ cells per 10 x 150 mm dish) were infected with each pooled library virus at low multiplicity of infection (MOI) achieving single viral integration and selected in medium containing puromycin (2 μg.ml^-1^) for 3 days. Post-puromycin selection cells were passaged for another 7 days to allow complete editing. Cells were trypsinized, pooled, counted and plated into 100 x 150 mm dishes (5 x 10^5^ cells per dish). 50 plates were infected with Ad-rtTA and 50 plates were infected with Ad-MAPL-flag adeno virus at 10 PFU/cell for 2 days. Medium was changed every 2-3 days. For Ad-rtTA, cells reached confluency in 1.5 week and for Ad-MAPL-flag, ∼ 3 weeks. Cells were harvested, trypsinized to single cells, pooled and mixed well. Approximately 20 x 10^6^ cells were used for isolating genomic DNA with High Pure PCR Template Preparation Kit (Roche) as described by the manufacturer. Library preparation for Next Generation Sequencing was done as described before (*53*). Briefly, the gRNA sequences were amplified from genomic DNA by PCR using Phusion HF DNA polymerase (ThermoFisher Scientific) using a 2-step amplification adding a unique 6-bp index per sample and sequencing adapter sequences. PCR products were purified using the High Pure PCR Product Purification Kit (Roche) and quantified using the Quant-iT™ PicoGreen™ dsDNA Assay Kit (ThermoFisher Scientific) before sequencing on a HiSeq2500 System (Illumina). Sequencing reads were mapped to the library using xcalibr and counts were then analyzed with MAGeCK (version 0.5.8) (*54*) using the Robust Rank Aggregation (RRA) algorithm to identify genes whose perturbation (knockout or overexpression) primarily enhanced fitness in the MAPL overexpressing group but not the control group. An sgRNA score was attributed to each sgRNA representing its relative abundance within each group. Guides with more than 1 mismatch and guides with less than 50 reads in rtTA library were filtered out. Reads of the remaining guides were normalised to the total (before filtering) sequencing library size for rtTA and MAPL overexpression libraries, and ratios of normalised averages (MAPL/rtTA) were calculated. The differential scores were calculated as log_2_(ratio of normalised averages), and ranked. Values with differential scores equalling zero were excluded from graphing. Calculations were computed using Rstudio version 2021.09.1, and graphs plotted using Prism10.

### Immunoisolation of lysosomes

U2OS cells stably expressing RFP-GAL3 and TMEM192-3xHA (*55*) were seeded overnight in 10 cm dishes (two dishes per condition, 2 x 10^6^ cells per dish). U2OS cells expressing RFP-GAL3 were also set up as a control not expressing TMEM192-3xHA. Lyso-IP U2OS cells were infected with 30PFU/cell adenovirus encoding DRING-FLAG or MAPL-FLAG. After six hours, virus was removed, and cells were washed three times in 1X PBS before addition of fresh media. Cells were left for a further 42 hours, before collection in PBS. Lyso-IP was undertaken as previously described (dx.doi.org**<u>/10.17504/protocols.io.bybjpskn</u>**) with modifications, as follows, based on (*56*). Cells were centrifuged once, then resuspended in 1 ml KPBS (136 mM KCl, 10 mM KH_2_PO_4_, 50 mM sucrose, pH 7.2). 2.5% input was isolated and resuspended in 1x Laemmli buffer. The remaining samples were lysed using 30 strokes in a Teflon-glass homogeniser. The homogenates were then centrifuged at 1000 *g* for 5 mins at 4°C. The supernatant was added to 100 μl of anti-HA magnetic beads (prepared by washing three times in KPBS) and placed on rotation for 20 minutes. The beads were then washed three times in KPBS, before elution in 2 x Laemmli buffer by heating to 95°C for 10 minutes. Samples were separated via SDS-PAGE (14% gel) before immunoblotting.

## FIGURE LEGENDS

**Figure S1.**
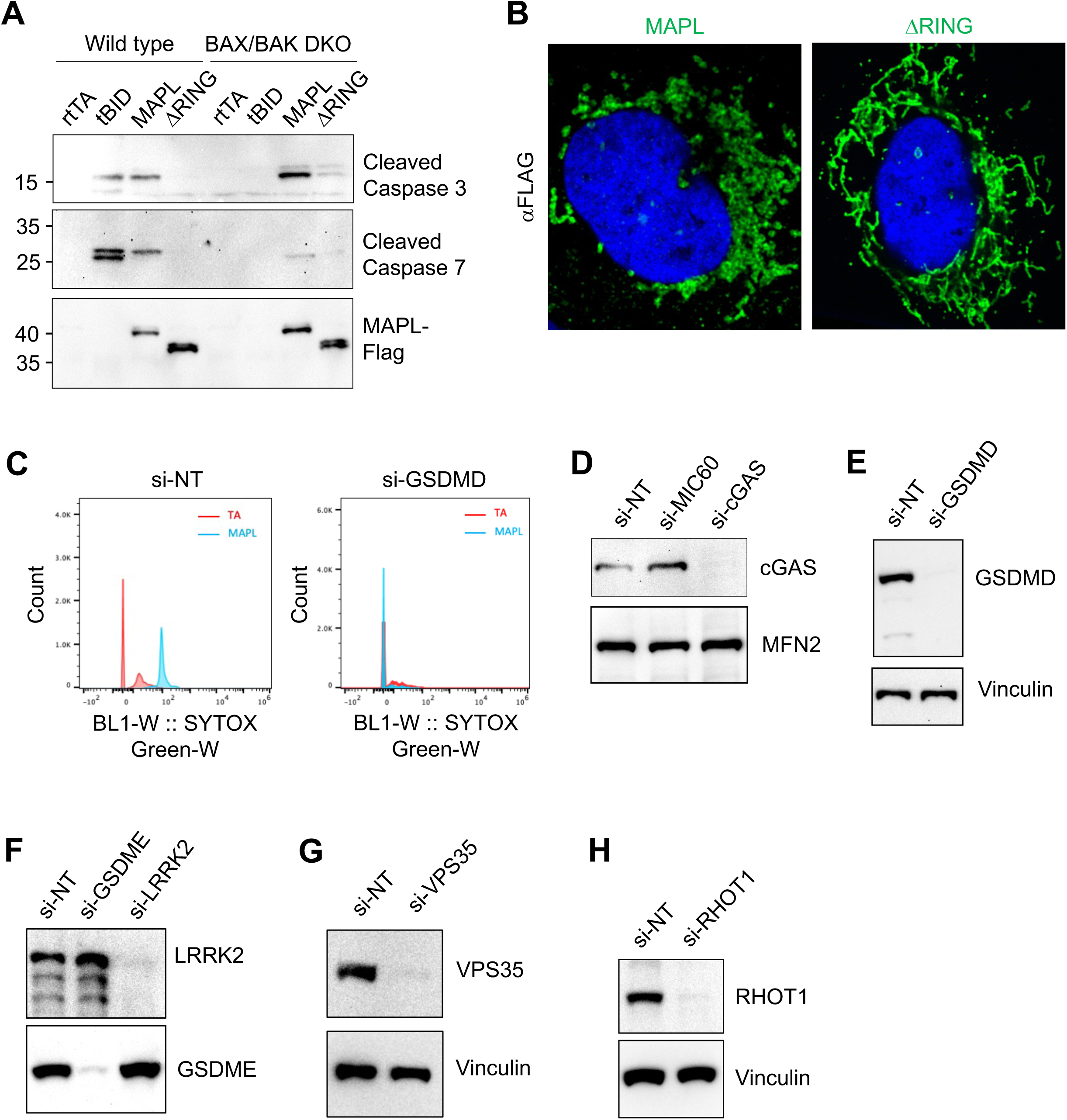
(A) MAPL promotes caspase cleavage in a BAX/BAK-independent manner (N=3). (B) Confirmation that adenovirus-mediated MAPL-FLAG and MAPL-ΔRING-FLAG localise to the mitochondrial network using anti-FLAG staining. (C) Flow cytometry revealed Gasdermin D (GSDMD) depletion prevent MAPL-induced SYTOX green uptake. (D-H) siRNA validations.

**Figure S2.**
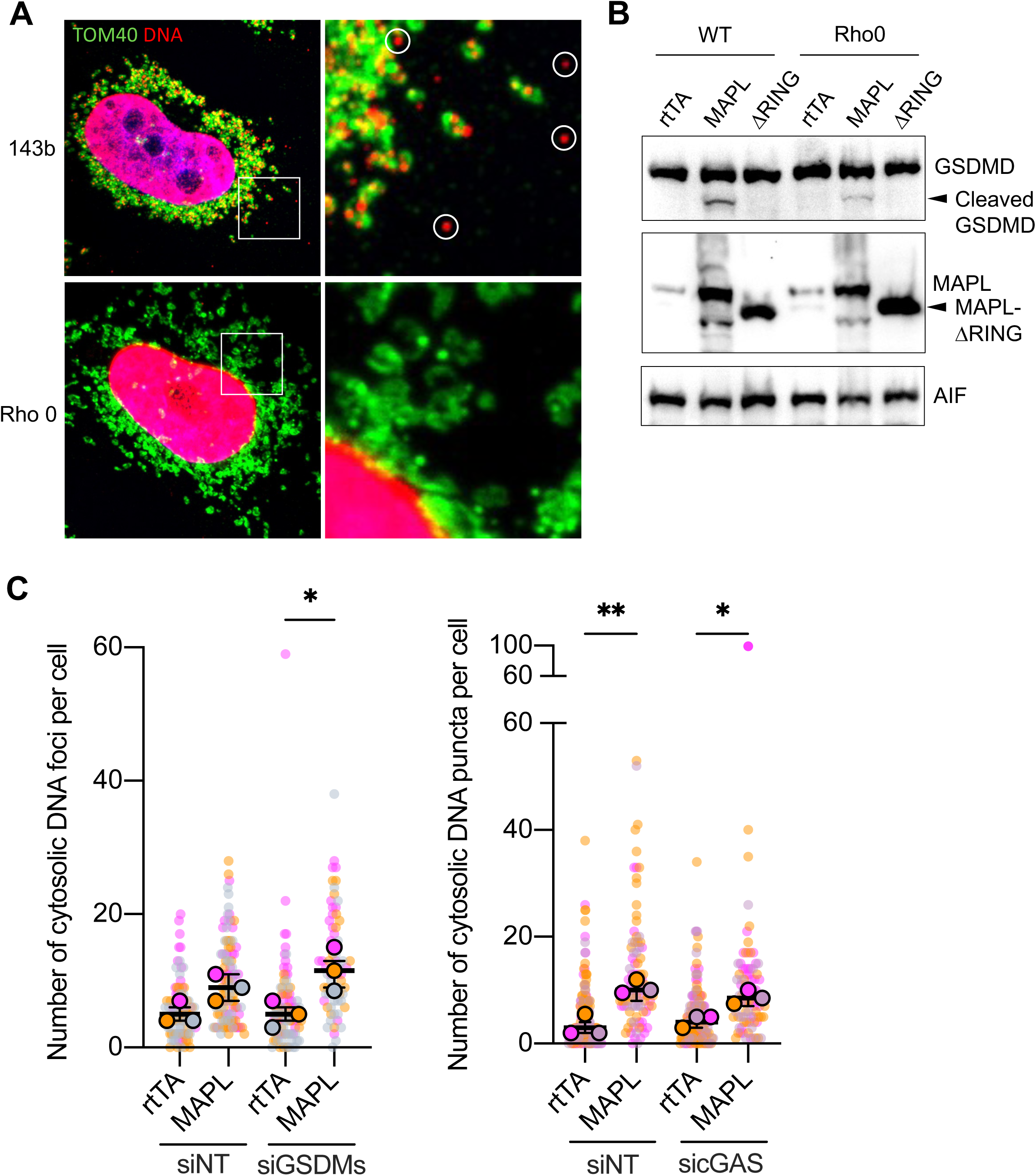
(A) No cytosolic DNA foci were observed in Rho0 cells after MAPL expression, whereas they were observed in parental 143b cells. (B) MAPL-induced Gasdermin D cleavage was inhibited in U2OS Rho0 cells generated by ethidium bromide treatment, compared to wild type (WT) cells. (C) cGAS or (D) Gasdermins (GSDMC/D/E) were not required for mtDNA release into cytosol after MAPL expression. Different colours represent independent repeats, with background circles showing corresponding values for individual cells. Values represent medians and error bars represent 95% CI. Comparisons were facilitated by One-way ANOVA. *P<0.05, **P<0.01.

**Figure S3.**
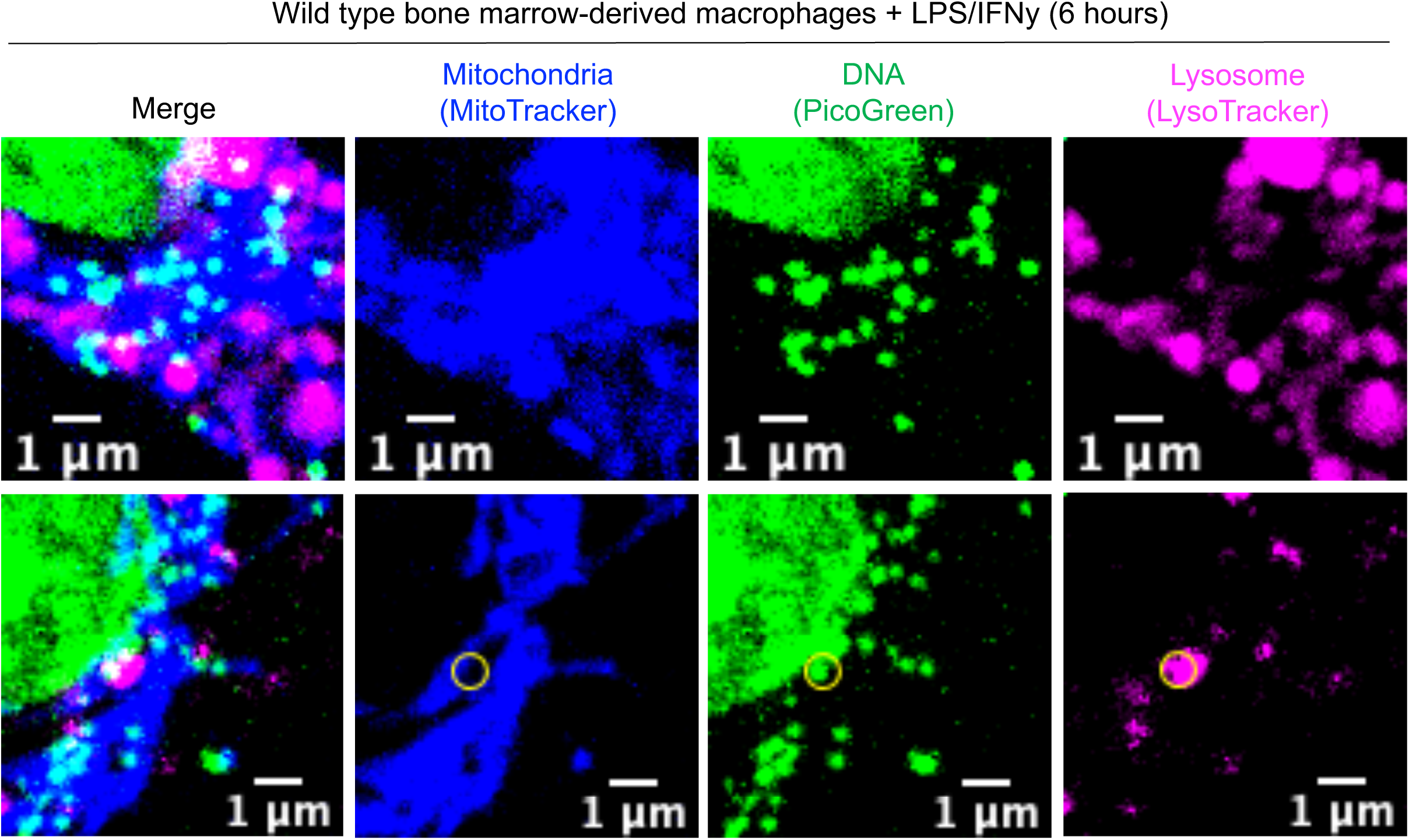
Bone marrow-derived macrophages were treated with lipopolysaccharide (LPS) and interferon-y (IFNy) for 6 hours. Circle shows DNA+ lysosome.

## REFERENCES AND NOTES

1. F. J. Bock, S. W. G. Tait, Mitochondria as multifaceted regulators of cell death. Nat Rev Mol Cell Biol 21, 85–100 (2020).

2. J. Prudent et al., MAPL SUMOylation of Drp1 Stabilizes an ER/Mitochondrial Platform Required for Cell Death. Molecular cell 59, 941–955 (2015).

3. V. Goyon et al., SUMOylation of ABCD3 restricts bile acid synthesis and regulates metabolic homeostasis. bioRxiv, 2022.2003.2003.482848 (2022).

4. P. S. Roque et al., Parvalbumin interneuron loss mediates repeated anesthesia-induced memory deficits in mice. J Clin Invest 133, (2023).

5. B. Zhang et al., GIDE is a mitochondrial E3 ubiquitin ligase that induces apoptosis and slows growth. Cell Research 18, 900–910 (2008).

6. E. Braschi, R. Zunino, H. M. McBride, MAPL is a new mitochondrial SUMO E3 ligase that regulates mitochondrial fission. EMBO reports 10, 748–754 (2009).

7. M. C. Wei et al., Proapoptotic BAX and BAK: a requisite gateway to mitochondrial dysfunction and death. Science 292, 727–730 (2001).

8. S. J. Korsmeyer et al., Pro-apoptotic cascade activates BID, which oligomerizes BAK or BAX into pores that result in the release of cytochrome c. Cell Death Differ 7, 1166–1173 (2000).

9. A. Weir, J. E. Vince, No longer married to inflammasome signaling: the diverse interacting pathways leading to pyroptotic cell death. Biochem J 479, 1083–1102 (2022).

10. P. Devant, J. C. Kagan, Molecular mechanisms of gasdermin D pore-forming activity. Nat Immunol, (2023).

11. S. D. Chauvin, W. A. Stinson, D. J. Platt, S. Poddar, J. J. Miner, Regulation of cGAS and STING signaling during inflammation and infection. J Biol Chem 299, 104866 (2023).

12. J. van der Beek, C. Jonker, R. van der Welle, N. Liv, J. Klumperman, CORVET, CHEVI and HOPS - multisubunit tethers of the endo-lysosomal system in health and disease. J Cell Sci 132, (2019).

13. S. M. Migliano, E. M. Wenzel, H. Stenmark, Biophysical and molecular mechanisms of ESCRT functions, and their implications for disease. Curr Opin Cell Biol 75, 102062 (2022).

14. A. Muraleedharan, B. Vanderperre, The Endo-lysosomal System in Parkinson’s Disease: Expanding the Horizon. J Mol Biol 435, 168140 (2023).

15. X. Zhang, X. C. Bai, Z. J. Chen, Structures and Mechanisms in the cGAS-STING Innate Immunity Pathway. Immunity 53, 43–53 (2020).

16. M. P. King, G. Attardi, Human cells lacking mtDNA: repopulation with exogenous mitochondria by complementation. Science 246, 500–503 (1989).

17. L. E. Newman, G. S. Shadel, Mitochondrial DNA Release in Innate Immune Signaling. Annu Rev Biochem, (2023).

18. K. McArthur et al., BAK/BAX macropores facilitate mitochondrial herniation and mtDNA efflux during apoptosis. Science 359, (2018).

19. N. Garcia, E. Chavez, Mitochondrial DNA fragments released through the permeability transition pore correspond to specific gene size. Life Sci 81, 1160–1166 (2007).

20. J. Kim et al., VDAC oligomers form mitochondrial pores to release mtDNA fragments and promote lupus-like disease. Science 366, 1531–1536 (2019).

21. H. Xian et al., Oxidized DNA fragments exit mitochondria via mPTP- and VDAC-dependent channels to activate NLRP3 inflammasome and interferon signaling. Immunity 55, 1370–1385 e1378 (2022).

22. C. G. Weindel et al., Mitochondrial ROS promotes susceptibility to infection via gasdermin D-mediated necroptosis. Cell 185, 3214–3231 e3223 (2022).

23. C. Rogers et al., Gasdermin pores permeabilize mitochondria to augment caspase-3 activation during apoptosis and inflammasome activation. Nat Commun 10, 1689 (2019).

24. D. V. Neel et al., Gasdermin-E mediates mitochondrial damage in axons and neurodegeneration. Neuron 111, 1222–1240 e1229 (2023).

25. H. Konaka et al., Secretion of mitochondrial DNA via exosomes promotes inflammation in Behcet’s syndrome. EMBO J, e112573 (2023).

26. T. Konig et al., MIROs and DRP1 drive mitochondrial-derived vesicle biogenesis and promote quality control. Nat Cell Biol 23, 1271–1286 (2021).

27. E. Braschi et al., Vps35 Mediates Vesicle Transport between the Mitochondria and Peroxisomes. Current Biology 20, 1310–1315 (2010).

28. V. Soubannier et al., A Vesicular Transport Pathway Shuttles Cargo from Mitochondria to Lysosomes. Current Biology 22, 135–141 (2012).

29. V. Soubannier, P. Rippstein, B. A. Kaufman, E. A. Shoubridge, H. M. McBride, Reconstitution of Mitochondria Derived Vesicle Formation Demonstrates Selective Enrichment of Oxidized Cargo. PloS one 7, (2012).

30. G.-L. McLelland, V. Soubannier, C. X. Chen, H. M. McBride, E. A. Fon, Parkin and PINK1 function in a vesicular trafficking pathway regulating mitochondrial quality control. Embo Journal 33, 282–295 (2014).

31. T. Eguchi et al., LRRK2 and its substrate Rab GTPases are sequentially targeted onto stressed lysosomes and maintain their homeostasis. Proc Natl Acad Sci U S A 115, E9115–E9124 (2018).

32. L. Bonet-Ponce et al., LRRK2 mediates tubulation and vesicle sorting from lysosomes. Sci Adv 6, (2020).

33. T. Kuwahara et al., Roles of lysosomotropic agents on LRRK2 activation and Rab10 phosphorylation. Neurobiol Dis 145, 105081 (2020).

34. S. Herbst et al., LRRK2 activation controls the repair of damaged endomembranes in macrophages. EMBO J 39, e104494 (2020).

35. N. Yadavalli, S. M. Ferguson, LRRK2 suppresses lysosome degradative activity in macrophages and microglia through MiT-TFE transcription factor inhibition. Proc Natl Acad Sci U S A 120, e2303789120 (2023).

36. J. Jia et al., Galectin-3 Coordinates a Cellular System for Lysosomal Repair and Removal. Dev Cell 52, 69–87 e68 (2020).

37. C. G. Weindel, L. M. Ellzey, E. L. Martinez, R. O. Watson, K. L. Patrick, Gasdermins gone wild: new roles for GSDMs in regulating cellular homeostasis. Trends Cell Biol, (2023).

38. G. A. Wyant et al., mTORC1 Activator SLC38A9 Is Required to Efflux Essential Amino Acids from Lysosomes and Use Protein as a Nutrient. Cell 171, 642–654 e612 (2017).

39. Q. Zhang et al., STING signaling sensing of DRP1-dependent mtDNA release in kupffer cells contributes to lipopolysaccharide-induced liver injury in mice. Redox Biol 54, 102367 (2022).

40. X. Zhan et al., LPS-induced mitochondrial DNA synthesis and release facilitate RAD50-dependent acute lung injury. Signal Transduct Target Ther 6, 103 (2021).

41. Z. Zhong et al., New mitochondrial DNA synthesis enables NLRP3 inflammasome activation. Nature 560, 198–203 (2018).

42. V. Zecchini et al., Fumarate induces vesicular release of mtDNA to drive innate immunity. Nature 615, 499–506 (2023).

43. M. F. Gulen et al., cGAS-STING drives ageing-related inflammation and neurodegeneration. Nature, (2023).

44. A. Sen et al., Mitochondrial membrane proteins and VPS35 orchestrate selective removal of mtDNA. Nat Commun 13, 6704 (2022).

45. J. Ding et al., Pore-forming activity and structural autoinhibition of the gasdermin family. Nature 535, 111–116 (2016).

46. X. Liu et al., Inflammasome-activated gasdermin D causes pyroptosis by forming membrane pores. Nature 535, 153–158 (2016).

47. N. Kumar et al., VPS13A and VPS13C are lipid transport proteins differentially localized at ER contact sites. J Cell Biol 217, 3625–3639 (2018).

48. L. Bonet-Ponce, M. R. Cookson, LRRK2 recruitment, activity, and function in organelles. FEBS J 289, 6871–6890 (2022).

49. T. Tang, B. Jian, Z. Liu, Transmembrane Protein 175, a Lysosomal Ion Channel Related to Parkinson’s Disease. Biomolecules 13, (2023).

50. D. Chatterjee, D. Krainc, Mechanisms of Glucocerebrosidase Dysfunction in Parkinson’s Disease. J Mol Biol 435, 168023 (2023).

51. Z. Harder, R. Zunino, H. McBride, Sumo1 conjugates mitochondrial substrates and participates in mitochondrial fission. Curr. Biol. 14, 340–345 (2004).

52. J. G. Doench et al., Optimized sgRNA design to maximize activity and minimize off-target effects of CRISPR-Cas9. Nat Biotechnol 34, 184–191 (2016).

53. S. Huang et al., MED12 controls the response to multiple cancer drugs through regulation of TGF-beta receptor signaling. Cell 151, 937–950 (2012).

54. W. Li et al., MAGeCK enables robust identification of essential genes from genome-scale CRISPR/Cas9 knockout screens. Genome Biol 15, 554 (2014).

55. M. Abu-Remaileh et al., Lysosomal metabolomics reveals V-ATPase- and mTOR-dependent regulation of amino acid efflux from lysosomes. Science 358, 807–813 (2017).

56. V. V. Eapen, S. Swarup, M. J. Hoyer, J. A. Paulo, J. W. Harper, Quantitative proteomics reveals the selectivity of ubiquitin-binding autophagy receptors in the turnover of damaged lysosomes by lysophagy. Elife 10, (2021).

